# Implementation of ddaE neuron growth mechanism in graph grammar replicates biological features

**DOI:** 10.64898/2026.07.27.741078

**Authors:** Matthew Hur, Patrick T. Hwu, Katherine L. Thompson-Peer, Eric D. Mjolsness

## Abstract

Dendrites develop branching patterns that are critical for their function, yet the mechanisms guiding arbor morphology remain incompletely understood, and quantitative models predicting how signals guide morphology remain limited. The ddaE neuron in *Drosophila* larvae is a proprioceptive sensory neuron with a characteristic asymmetric dendrite arbor that exhibits posterior-biased branching. We developed a computational model using Dynamical Graph Grammar (DGG) to simulate ddaE dendrite development as a graph-based dynamical system, using a single morphogen gradient to establish arbor architecture. Our simulations of ddaE dendrites, guided by the spatial gradient of the Teneurin-m (Tenm) morphogen combined with resource constraints and self-avoidance rules, accurately recapitulate the morphological features of biological ddaE neurons, including primary branch orientation, posterior bias, branch tree distributions, and branch length statistics. We find that the response to a single morphogen gradient is sufficient to guide the computerized dendritic arbor. Null model analyses demonstrate that simulated arbors exhibit non-random spatial and topological organization consistent with biological constraints. Our results demonstrate that rules based on a single morphogen gradient are sufficient to generate complex asymmetric dendritic patterns and provide a validated computational framework for testing perturbations *in silico*.

**SIGNIFICANCE:** We develop and simulate a minimal computational model of dendritic arbor morphogenesis based on a single morphogen gradient. Asymmetric dendrite arbors, such as the ddaE proprioceptive neuron in Drosophila larvae, have not previously been computationally modeled. Using the Dynamical Graph Grammar framework, we create a mathematical model from 17 dynamical rules governing changes in arbor structure, spatial position, morphogen-directed growth, and the local dynamic state of each tip. Each tip switches among three states: growth/pause/shrinkage, while tips that encounter another branch additionally enter a retraction state. Using a large simulation sample size, we characterize our model system’s generative outputs and verify that they match imaged biological dendrites across the majority of morphological statistics, including posterior branch bias, branch degree distributions, branch number, and dendrite length. We demonstrate that efficient spacing is guided by the orientation of branch junctions. We make our simulation publicly available to function with high-throughput investigation of gene-to-phenotype relationships in dendrite development.

## INTRODUCTION

Dendrite morphology determines neuronal function by defining the spatial domain of synaptic integration and, in sensory neurons, the receptive field for stimulus detection. Understanding how stereotyped dendritic arbor patterns emerge from molecular and biophysical processes remains a fundamental challenge in cellular neuroscience. The transformation of molecular cues like morphogen gradients into complex three-dimensional branching structures requires quantitative frameworks that can bridge scales from molecular signaling to tissue-level organization. Computational modeling provides a powerful approach to test whether proposed molecular mechanisms are sufficient to generate observed morphologies and to make quantitative predictions for experimental validation.

In the *Drosophila* larval peripheral nervous system (PNS), a pair of class I dendritic arborization (da) neurons, named ddaE and ddaD, serve as key proprioceptors that detect body curvature during forward and reverse locomotion, respectively. ddaD and ddaE neurons exhibit highly stereotyped and reproducible asymmetric morphologies, making them an ideal system for quantitative modeling. ddaE dendrites extend towards the anterior end of the larvae and primarily detect mechanical curvature during forward locomotion, while ddaD dendrites extend towards the posterior end of the larvae and detect mechanical curvature during backward locomotion (1, 2). Thus the morphology and architecture of these dendrite arbors directly determine the function of the neurons, making them excellent model systems for understanding structure-function relationships. Yet despite ddaE’s and ddaD’s robust patterning, we currently lack a computational model that recapitulates how this stereotyped development is achieved. Developing such a tool will provide new opportunities to identify factors that contribute to dendrite outgrowth.

This paper focuses on ddaE neurons, which exhibit a stereotyped and highly reproducible morphology that makes them amenable to computational modeling. ddaE neurons have 2-3 primary branches that extend from the cell body: one long dorsal branch extending upward, and one (or two) slightly shorter branches extending in the ventral-anterior direction (3). The dorsal branches exhibit strong posterior bias in secondary and higher-order branching, forming a distinctive comb-like morphology, with secondary branches extending posteriorly. The dorsal arbor is also consistently larger than the ventral-lateral arbor. This posterior bias is guided by a morphogen gradient of the cell adhesion molecule Teneurin-m (Ten-m) (4). Ten-m is expressed by class I da neurons and by epithelial cells anterior to the ddaE cell body in a dorsoventral stripe. Ten-m acts as a homophilic adhesion protein, inhibiting anterior ddaE dendrite outgrowth by mediating adhesion between the dendrite membrane of the da neuron and Ten-m expressing epithelial cells (4).The ventral part of the ddaE arbor is angled with respect to the dorsolateral and anteroposterior axes (~30° downward) (3). Thus, ddaE development presents a well-characterized system in which a defined molecular gradient creates an asymmetric dendritic field.

Despite identification of some key guidance molecules, fundamental biophysical questions remain unanswered. For example, is absolute morphogen concentration or spatial gradient the critical parameter for guiding branch growth? Are additional molecular pathways required beyond what has been identified, or is a single cue sufficient to specify the complete arbor pattern? What biological constraints, such as resource availability or self-avoidance mechanisms, shape final arbor morphology? Answering these questions requires a computational framework that can: represent dendrites as dynamical branching structures; incorporate biophysical rules for growth, branching, and retraction; respond to spatial molecular cues; and be quantitatively validated against experimental data.

Several computational approaches have been developed for modeling dendrite morphogenesis, each with distinct strengths and limitations. A model that connects scaling of involved modules, e.g. the glomerulus and the olfactory bulb, provides an explanation of sensory function from scaling of basic phenotype (5). Since that modeling lacks exogenous signaling and relies on a few simple mechanisms, the system complexity is non-trivial to observe from simulation. Previous work has modeled Drosophila class IV da neurons, which exhibit space-filling arbors with different patterning principles than class I neurons (6, 7), and symmetric class I neurons (8), but asymmetric class I neurons like ddaE have not been computationally modeled. A key challenge across all approaches is quantitative validation: models must not only quantitatively resemble biological morphology but also match statistical distributions of branch angles, lengths, degrees, and spatial organization.

In this paper, we address this gap by developing a computational model of ddaE dendrite arbor development, using the Dynamical Graph Grammar (DGG) modeling language package, Plenum (9), written in the Mathematica computer algebra system. The DGG language (10) provides a powerful framework for modeling biological morphogenesis, as it represents morphological structures as dynamical graphs and specifies growth through formal rewriting rules. The DGG approach is well-suited for dendrite modeling because: dendrites naturally form tree graphs with branches as edges and branch points as nodes; discrete events like branch addition, retraction, or bifurcation map directly to graph rewriting operations; stochastic dynamics emerge naturally from rule firing rates; and spatial embedding allows incorporation of molecular gradients and other geometric constraints. DGG has previously been used to simulate synaptic spine head dynamics and internal actin biophysical dynamics (11), as well as cortical microtubule remodeling (12), but has not previously been applied to whole-cell dendrite morphogenesis.

Our model incorporates three core biophysical components. First, morphogen guided growth: branch elongation velocity is determined by the spatial gradient (finite difference) of Ten-m concentration rather than an absolute level, creating spatial memory in the growth process. Second, resource allocation constraints: a resource allocation function limits the total arbor size and implements the observed dorsal-ventral growth bias (the dorsal arbor grows faster than the ventral). Third, self-avoidance: dendrite branches avoid crossing through a mechanism to occupy and control space. We implement these components as graph rewriting rules in the Plenum DGG software package, specify model parameters based on experimental measurements where available, estimate remaining parameters where necessary, and run ensemble simulations to generate populations of synthetic arbors.

Here, we present a calibrated and validated computational model of ddaE dendritic development in the DGG modeling language. We systematically compare our DGG simulations against biological ddaE neurons at multiple levels of quantitative analysis, including distributions of branch numbers, lengths, degrees, and posterior bias to establish model accuracy and predictive capability. We further assess model specificity against null models with randomized branch arrangements and spatial distributions. Together, this multi-level validation framework establishes model accuracy and provides a quantitative foundation for future *in silico* exploration of molecular perturbations, parameter sensitivities, and alternative mechanistic hypotheses underlying asymmetric dendrite development, regeneration, and the relationship between dendrite structure and sensory function.

## MATERIALS AND METHODS

### Biological image collection

Drosophila melanogaster were raised on a standard cornmeal yeast diet at 23° C. The following fly stocks were used: Canton-S (wild-type), and ppk-gal4; ppk-cd4-tdgfpÎb (13). Uninjured class I ddaE neurons were imaged 96 hrs after egg lay (AEL) on a Zeiss LSM 700 confocal microscope. Larvae were mounted dorsal side up in glycerol. Z-stacks spanning the full depth of the neuron were acquired in 1 *μ*m increments using a 20x objective with a 488nm excitation. Maximum intensity projections were generated in Fiji/ImageJ (14) for downstream analysis. A total of 27 neurons and 22 larvae were imaged and analyzed.

### Image Analysis

Dendrite arbors were traced and analyzed from maximum intensity projections using the Simple Neurite Tracer (SNT) plugin (14) in Fiji/ImageJ. Neurons were traced by one independent tracer. From each traced arbor, the following morphological features were extracted: total tip number, total length, branch degree (primary, secondary, tertiary, quaternary), average and total branch length per category, and anterior-posterior counts. Branches were classified as dorsal (top arbor) or ventral-lateral (bottom arbor) based on their origin from the dorsal or ventral-lateral primary dendrite, respectively. Branch degree is defined as the number of branch points away from the soma, with the primary branch attached to the soma (not to be confused with the definition of degree in graph theory). Within each arbor, branches were classified as posterior or anterior based on their direction of extension relative to the anteroposterior body axis of the animal. All successfully imaged neurons were included in the analysis.

### Computational model overview

The dendrite is represented as a graph by a set of “Node[…]” objects, whose graph relationships are represented by shared parameter values. The parameters include the ID, spatial coordinate, previous node ID, next node ID, branching node ID, growth state, node degree, node arbor side, node arbor top vs. bottom, and x and y grid IDs that belong to a partition of the coordinate axes. A branch tip as a Node object possesses a null pointer for its “next node ID”. The initial condition for all simulations is five nodes directly upwards from the soma and three nodes with a 30° ventro-posterior bias for the bottom arbor.

We use a modified algorithm of the Stochastic Simulation Algorithm (SSA) that allows for parameterized objects and propensities as functions of the parameters (9, 15). The objects update according to user-specified rules written in the Plenum modeling language package in the Mathematica computer algebra programming language. Simulation iterations occur until the 1200 minute mark is reached and take roughly an hour of wall clock time. 100 such simulation runs comprise our set for analysis.

### DGG rule implementation

The following rules govern the stochastic evolution of the dendritic arbor graph, including branch elongation, branching, node retraction, growth state transitions, and self-avoidance.

#### Branch elongation

The branch elongation rule governs the extension of dendrite branch tips. In the “with” clauses we have converted the rate constant to a function of growth rate, resource limitation, and Ten-m level finite difference. *R* (*θ*) is a 2 × 2 rotation matrix with angle *θ*. The branch elongation rule fires at a propensity that is controlled by a resource constraint scaling function (i.e. Eq. 4), the Ten-m morphogen gradient as input to a logistic function (i.e. Eq. 3), and the actual growth rate (i.e *V*_*G*_ from Table I) with a change of units of velocity of *μ*m/s to nodes/s (i.e.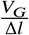). For the dorsal (top) arbor, (i.e.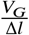) is scaled by *r*_dorsal_ to model the greater amount of resources supplied to the dorsal arbor. To mathematically represent the Boolean choice of dorsal or ventral arbor for *r*_dorsal_, we compare *b*_dv_, the categorical input of dorsal or ventral arbor to the actual categories by the Kronecker delta function defined in Eq. 1.

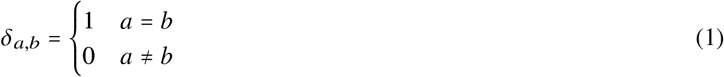

In addition, a factor of two is multiplied inside the “with” clause of node addition rules such as Eq. 2 for the rule to fire sequentially with one other self-avoidance checking rule (Eq. 8).

#### Branch Elongation

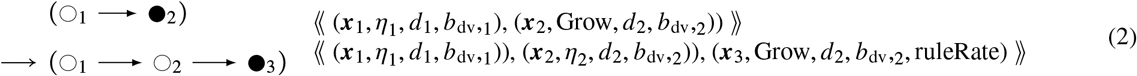

Here *η* represents the growth state of the node, which matters for end nodes. The “;” represents sequential execution of two successive parameter assignments, so that *θ*_2_ is sampled before use in setting ***x***_3_. The sampling of *θ*_2_ is centered around zero as its mean, with a standard deviation *σ*_*θ*_ taking into account the persistence length *L*_*p*_. As shown in the “where” clause in Eq. (2), *η* represents the growth state (one of Grow, Pause, or Shrink) attached to a node object. This paradigm was empirically observed and characterized in previous studies from which we use their growth and shrinkage velocities as well as the three-state transition rates (6, 7). *α* is a counter for the branch degree of the node. *σ*, is a class for either the top or bottom arbor. There are other nodal object parameters, suppressed here for readability. Node type symbols and represent interior and end segments of a dendrite branch.

We apply resource constraints using a function of three parameters: the simulation state’s arbor size (*α*), which is separated into top arbor size and bottom arbor size, the branch degree of the node location (*d*), and a binary variable (*b*_*dv*_) for whether the node is in the dorsal (top) or ventral (bottom) arbor. *α* is the scaled input to the logistic function, where a large arbor size input to the logistic function leads to a saturating value output close to 1. *S* is a fixed arbor size value that scales *α* to a value close to 1. We horizontally flip the logistic function by negating the input. It results as a decreasing function from 1 to 0, to lessen propensity with which the arbor growth rules (Eq. 2, Eq. 5) fire as the arbor size grows. In this way, resource limitations are implemented in the form of feedback from arbor growth into smaller growth rule propensities. The degree *d* also scales the growth propensities as part of the resource function. Branches higher degrees away from the cell body grow slower to model the resource and transport limitations. The factor of 2 in Eq. 4 below defining the resource function serves to equate the effective starting growth rate (where *α* = 0) of an arbor with a plausible value provided in Table I. The function “resourceFunction” in the “with” clause of Eq. 2 is a saturating logistic function, i.e.

**Table I:** Table of parameters used in simulation of ddaE da class I neuron.

| Parameter | Value | Description |
| --- | --- | --- |
| $\Delta l$ | 12.5 $\mu\text{m}$ | Edge length between two consecutive nodes |
| $V_G$ | 1.61 $\mu\text{m}/\text{min}$ | Growth velocity in growth state (7) |
| $V_S$ | -1.53 $\mu\text{m}/\text{min}$ | Growth velocity in shrink state (7) |
| $V_{S,\text{contact}}$ | -1.9 $\mu\text{m}/\text{min}$ | Self-avoidance shrink velocity (6) |
| $r_{\text{branch}}$ | 0.0016 1/min 1/$\mu\text{m}$ | Branching rate (6) |
| $k_{GP}$ | 0.784 1/min | Growth to pause state transition rate (7) |
| $k_{GS}$ | 0.640 1/min | Growth to shrink state transition rate (7) |
| $k_{PG}$ | 0.335 1/min | Pause to growth state transition rate (7) |
| $k_{PS}$ | 0.314 1/min | Pause to shrink state transition rate (7) |
| $k_{SG}$ | 0.598 1/min | Shrink to growth state transition rate (7) |
| $k_{SP}$ | 0.946 1/min | Shrink to pause state transition rate (7) |
| $L_p$ | 900 $\mu\text{m}$ | Branch persistence length |
| $r_B$ | 0.0016 1/min 1/$\mu\text{m}$ | Branching rate (6) |
| $T_{\text{ten-m}}$ | 0.0035 | Logistic function temperature for Ten-m gradient response<sup>a</sup> |
| $r_{\text{dorsal}}$ | 1.3 | Ratio and multiplier of dorsal to ventral arbor resource allocation |

<sup>a</sup>The gradient finite difference is normalized to the maximum value before multiplying by inverse temperature.

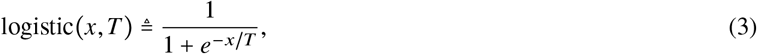

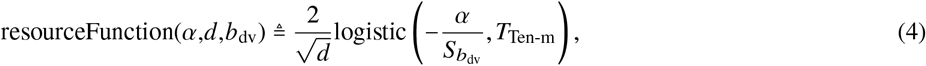

that takes as input the size of the top or bottom arbor, which is stored in either the top or bottom node count objects sampled concomitantly with the node objects part of the dendrite graph. In Eq. 2 and 5, the logistic function following “resourceFunction” is a response function to the finite difference of the Ten-m gradient the stability of which can be controlled by the temperature parameter (Table I). The growth velocity of branch in Table I, *V*_*G*_ is multiplied by the asymmetric resource allocation that is higher in the dorsal arbor.

#### Branching, retraction, and growth state transitions

##### Branching

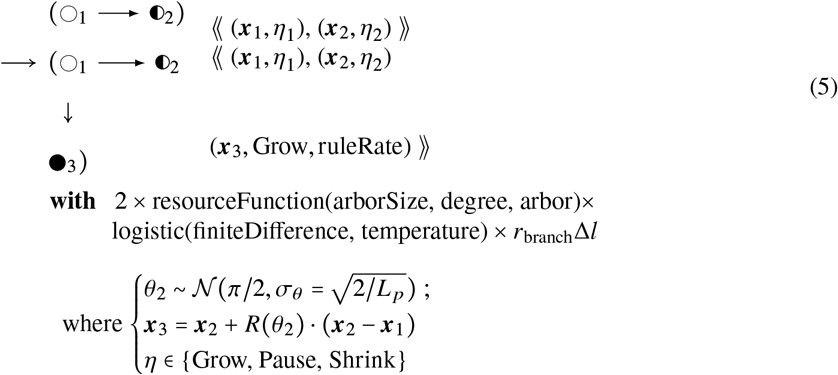

The parameter *r*_branch_, used in Eq. 5 and provided in Table I is the rate per unit length that a branching event occurs. This parameter is converted to branching rate per node by multiplying by Δ*l*. The rest of the propensity “with” clause is lifted from Eq. 2, which is explained earlier in this section.

##### Node retraction

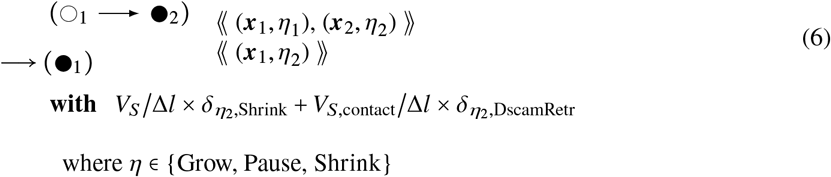

The parameters *V*_*S*_ and *V*_*S*,contact_ are provided in Table I as retraction rates, with *V*_*S*_ being the default retraction rate for a tip in a retraction state and *V*_*S*,contact_ the self-avoidance retraction rate. Self-avoidance is triggered for a tip that collides with a branch and retracts to the nearest arbor junction. The choice between retraction rates is represented mathematically by a Kronecker delta function *δ* (Eq. 1) that compares the *η*_2_ tip state parameter with the respective state, i.e. either shrink or Dscam retraction.

**<H>Growth state transition**

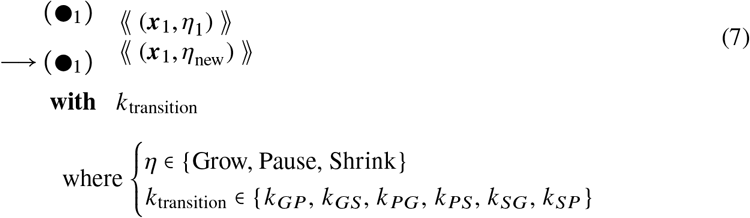

{*G P S*}

The parameter for the state of the branch tip *η*_1_ in Eq. 7 is mapped to the propensity variable *k*_transition_, which depends on *η* as a transition between two states from {*G*, *P*, *S*}. The single letter symbols represent states as follows: *G* →Grow, *P* → Pause, and *S* → Shrink. *k*_transition_ selects two of the possible values for *η* and represents a transition from *η*_1_ to *η*_new_ where the transition subscript, “transition” = *η*_1_*η*_new_.

#### Self-avoidance checking and implementation

In addition, to check for self-avoidance (dendrite branch cross-overs) with a method that is compatible with Dynamical Graph Grammars, we add onto object three’s parameters the rate inside the “with” clause that was multiplied by two to account for the subsequent self-avoidance check rule. In this way, before another rule firing adds another node, a different rule needs to return the number of parameters back to the number present in the left-hand-side of Eq. 2 after checking for cross-overs.

The rule for self-avoidance checking (Eq. 8) uses sub-grammar calls to implement an OR operation that chooses one of two rules, depending on whether there is any third node (subscript 3) in the same grid location as the tip node (subscript 2). If there exists a third node in the grid location of the tip node, the tip node receives a tip state of “DscamRetr” in order to retract back to the nearest junction, for which case only the node retraction rule for DscamRetr fires in Eq. 8. The sampled object that allows for the OR operation is the flag ⋁ that gets sampled and re-created as ⋀ after either of the two sub-grammars operating on the dendrite tree fires. Then, ⋀ is re-created as ⋁ to reset the ability of Eq. 8‘s sub-grammars to fire in the final sub-grammar call of Eq. 8.

The sub-grammar calls are necessary in this case, in order to mandate through rules the deterministic checking after every node addition for collisions of tips with branches. The OR operation, which needs sub-grammar calls for a single rule, lets exist two cases: either the checking notices a collision or the checking allows continual growth.

The self-avoidance checking rule of Eq. 8 fires at a rate that was intentionally set to two times the desired firing rate in Eq. 2 and Eq. 5. This rate was stored in the tip object, as an additional parameter, after the growth rule had created it. The additional parameter prevents the other tip sampling rules from pattern-matching for that tip until the self-avoidance checking rule sets the number of parameters without the valued parameter symbolized as “ruleRate” in Eqs. 2 5, and 8.

We create DGG rules for self-avoidance using the following notation: ⋀, ⋁, and 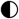. The symbol denotes a flag object that is “active” and ⋁ denotes a flag object that is “inactive” with a Boolean 1 or 0 stored inside the brackets of “flag[…]”. The symbol 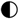 represents either ○ or 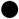. The diagrammatic rule for self-avoidance checking is shown below:

##### Self-avoidance Checking

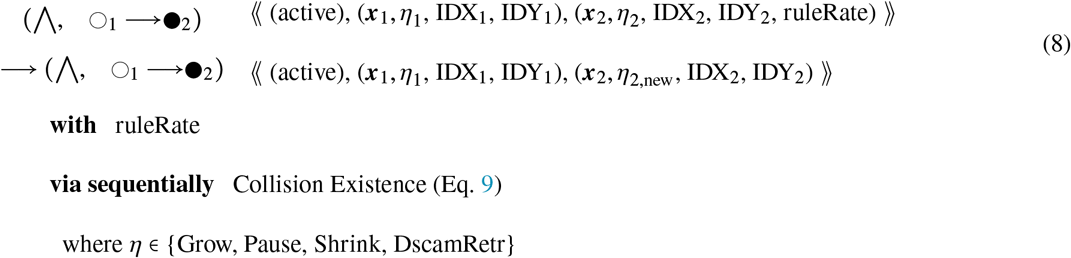

##### Collision Existence

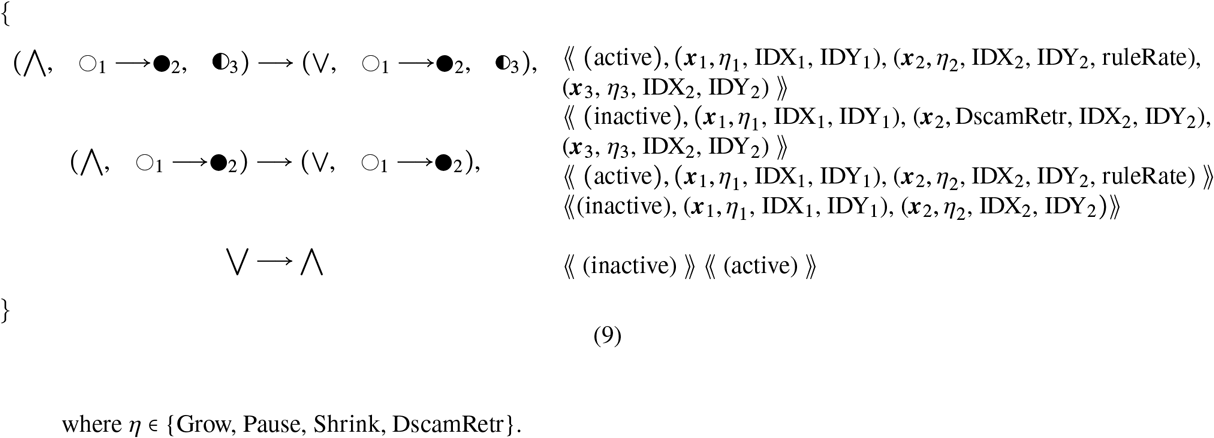

The super-grammar (Eq. 8) calls the sub-grammar (Eq. 9), whose rules are applied one after another in the listed order, using the “via sequentially” keyword phrase, whose semantics is similar to the “via” dynamical grammar rule keyword introducing fast subgrammars (16), but with a local sequentialization of “via” subrules that could be implemented without use of the “sequentially” keyword, by an extra support object with a discrete step parameter similar to a program counter running from 0 to 3, the number of subrules. This subrule mechanism is a standard feature of the Plenum implentation of Dynamical Grammars (9).

To fire, the outer LHS must find a match within the pool of objects, which includes the end node 2. This end node must have one more parameter, i.e. the propensity firing rate (“ruleRate”) for the rule. In the RHS, three more rules are specified as sub-grammar rules. They alter a “flag[…]” object containing a single Boolean parameter, which is shown in Eq. 8 as active, i.e. ⋀, or inactive, i.e. ⋁. The inclusion of this flag ensures that only one of the two rules that alter the growth state of the end node 2 fires within the RHS. The first of the two rules checks for an existence of a third node possessing the (IDX_2_, IDY_2_) parameter and sets the growth state to self-avoidance for end node 2. The second of the two rules removes the extra firing rate parameter stored in end node 2 to enable it for further graph-topological change. The parameter *η*_2_ corresponds to an input “Growth” state that can change by sub-grammar call, to either “DscamRetr” or remain at “Growth” as the final state represented by the parameter *η*_2,new_.

The raw code for this self-avoidance checking rule is provided in Fig. II. We implement a deterministic self-avoidance mechanism as a pair of DGG rules. The method handles the immediate retraction following a dendrite branch cross-over event. The self-avoidance mechanism of ddaE arbors (17–19) is likely based on the heterophilic repulsion molecule, Dscam, and is implemented with our new DGG method. This method extends a pool of simulation dendrite objects spanning a parameterized spatial grid with one new object for each grid spot. Instead of programming a combinatorial check for all branch tips against all pairs of connected branch nodes, if a branch tip elongates into an occupied grid spot, the self-avoidance mechanism triggers. Once self-avoidance is triggered, the branch retracts at a velocity faster than the shrinking state velocity back to the previous branch point.

**Figure 1:**
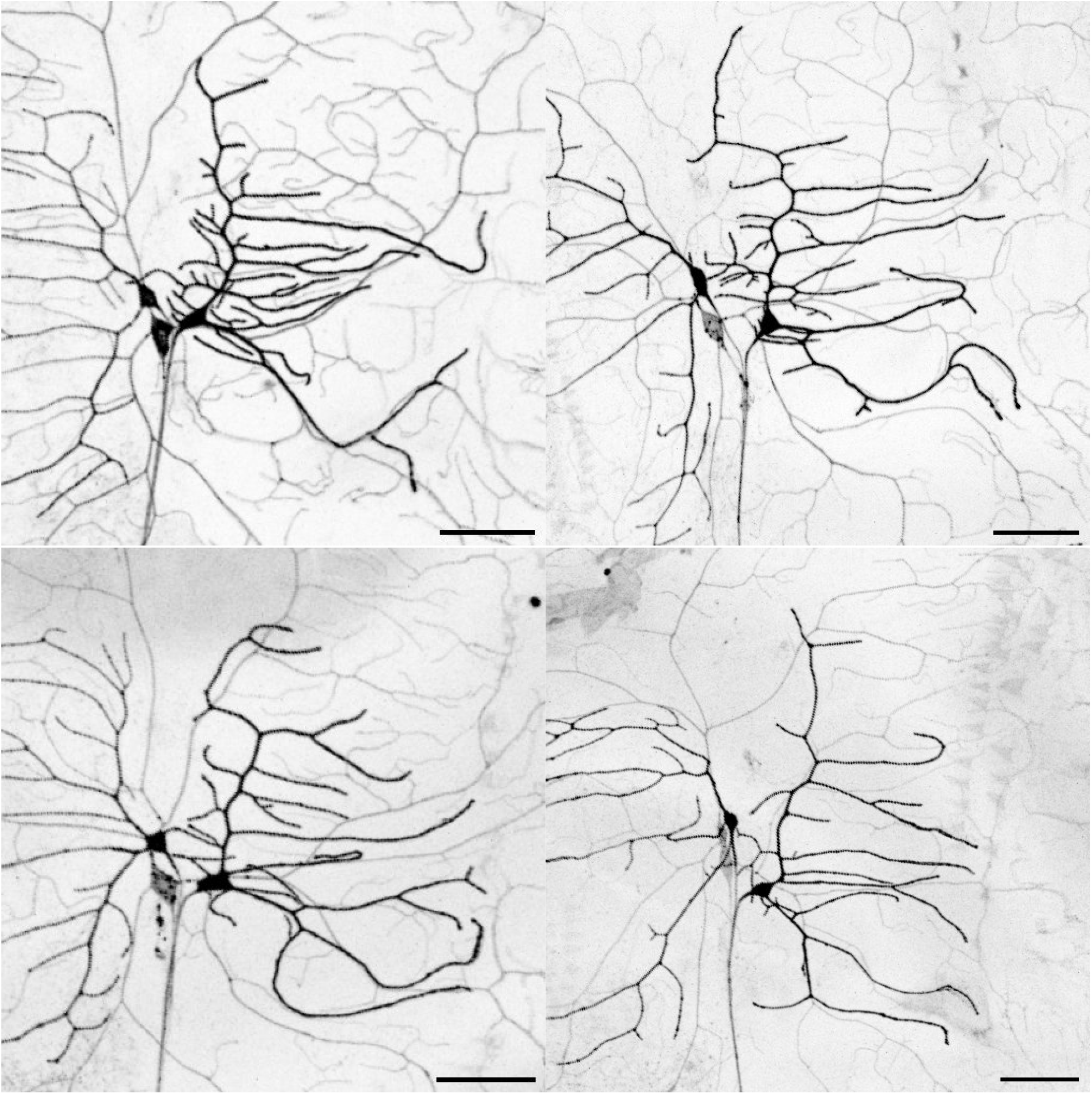
Four example images of ddaE biological dendrites. Dendrites are dark black projections, cell bodies are near the center, and axons extend downwards. ddaD dendrites grow left beyond the field of view (anterior) and ddaE dendrites (the subject of this study) grow right (posterior). Class IV ddaC neurons are visible in light gray in the background. 50*μ*m scale bars.

**Figure 2:**
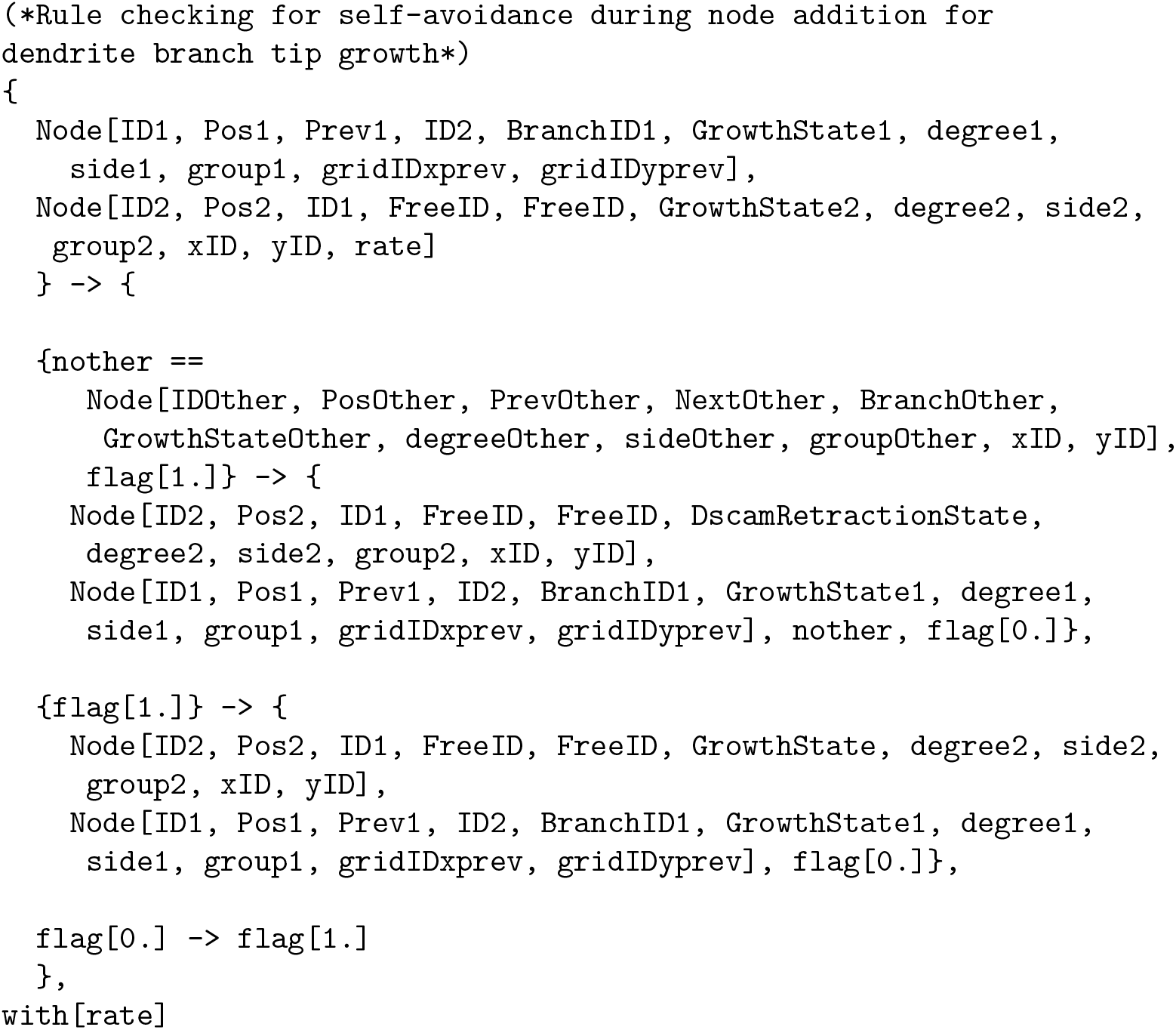
Actual Plenum rule text of rule of Eq. (8), firing by design after a separate rule-firing instance of Eq. (2). “flag[1.]” is an object initialized in the pool before the first iteration of the simulation. Node objects containing ID1 and ID2 are the penultimate and end nodes of a dendrite branch. “nother” is a potentially existing object in the same grid spot, localized as (xID, yID) from Node ID2. If that node does exist, then ID2’s growth state is set to “DscamRetractionState” for which different rules fire for self-avoidance retraction. “GrowthState” is a variable stored with a numerical value for pattern-matching. In this particular instance of a Plenum DGG rule with DGG subrules, the output graph is provided by either the first or second subrule (at least one of which will execute) rather than, as would be more common, being listed explicitly on the right hand side of the main rule before any of the subrules are listed.

This checking step requires a follow-up rule that acts immediately after each node addition. By taking advantage of a rule-attached clause named “via” in the declarative modeling language of DGGs, supported in Plenum, the DGG grammar chains a rule with a sub-rule together in rule firing sequence. We use rules specified with “via”, together with parameterized object-matching, to implement self-avoidance, checking for the presence of a non-immediately connected node in the same grid spot as the proposed node. This rule’s raw code is shown in Fig. II.

### Ten-m morphogen gradient

A piecewise linear ramp function of time for the Ten-m gradient initiates from *t*_start_ and ends at *t*_end_ over a temporal duration *t*_window_. This interval is represented by the difference of Heaviside step functions, i.e *H*(*t* −*t*_start_) −*H*(*t* −*t*_end_). For when *t* > *t*_end_ without a ramp scaling multiplicative term, *H* (*t* − *t*_end_) adds to the interval with the scaling multiplied as a total to be multiplied to the spatially multivariate Gaussian.

Eq. 10 represents the concentration of the Ten-m morphogen molecule across the spatial locations where the dendrite grows. The Gaussian assumes a morphogen peak coordinate location of 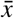 and morphogen concentration standard deviation of *σ*. Location and width of the Ten-m gradient was estimated from (4). The gradient assumes a Gaussian form as would be expected to result from a diffusion process (not explicitly modeled), i.e. approximately

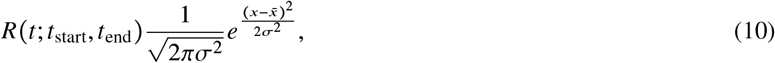

where *R*(*t*; *t*_start_, *t*_end_) is the bounded ramp function

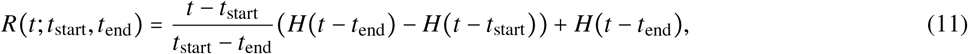

where *H*(*t*) is the Heaviside step function, *t*_window_ = *t*_end_ − *t*_start_ is the time for source to be fully active, 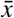 is the horizontal mean of the gradient, and *σ* the horizontal standard deviation. The Ten-m gradient mean is − 40 *μm* and standard deviation 45 *μm* where the cell body is located at coordinate {0, 0}.

The temperature for the response to the finite difference of the Ten-m gradient, *T*_ten-m_ was hand-tuned to match the biological dendrite images’ anteroposterior asymmetry distribution. The finite difference is computed as a difference between the proposed growth location and the current end: 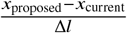. Finite differences can create memory within the morphological growth despite DGG rules generating dendritic arbors based on the Stochastic Simulation Algorithm as a Markov chain. The gradient as a derivative determines the degree of growth which leads to activation where it is changing and not at one of the ends of a saturating logistic function. A finite difference of the morphogen gradient can cause associative learning of the important region of the gradient where it is changing.

### Resource allocation between top and bottom arbors

For our resource constraint, we bias the growth of the top arbor by allocating to it 1.3 × the resource for the bottom arbor. Additional objects are stored in the object pool for the total number of top or bottom da node objects, updated at each node addition or deletion rule firing. Also, we assume that growth velocity of dendrite tips in the top arbor is 1.3 × faster than the bottom arbor because of the higher level of growth resource molecules such as actin.

### Null model mathematics

We use several natural null models to understand our simulation data. The first or “dartboard” null model comprises a probability distribution on sets of points imagined as darts thrown at a rectangular dartboard, representing features of the discretized dendritic tree including branch points or junctions, branch termination points, and/or internal nodes. The probability model is an independent uniform distribution on the rectangle for each such point. There is a separate rectangle for the top and bottom portions of the dendritic tree, each constrained to be within the bounding box of the corresponding region of a simulated dendrite arbor. The resulting “dartboard” null model for randomly spaced points can be compared to the feature points of a simulated arbor grown by the DGG rules.

A key statistic to compute for each use of the dartboard null model is a within-rectangle histogram of distances, expressible mathematically as

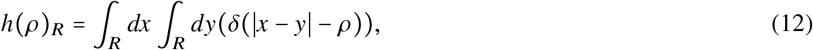

where *h* (*ρ*)_*R*_ is a histogram of the distances of radii centered at each relevant node in the simulation tree; *x* and *y* are two sampled coordinates; *ρ* is the radius of the search circle, and *δ* is the Dirac delta function.

A second “Galton-Watson tree” null model is named after a standard model of descent in which each node can have zero, one, or two unordered children nodes with constant probabilities, independent for each node, that add up to unity. One sometimes conditions this distribution on the total number of nodes in the tree before all final nodes have zero children each. *p*_branch_ represents the chance that a node has two outgoing edges, and *p*_leaf_ represents the pooled fraction of leaf nodes as the terminal leaf probability of having zero outgoing edges. Then, the probability of having a single outgoing edges is 1 − *p*_branch_ −*p*_leaf_. A random Galton-Watson tree, where the number of children at a node is randomly sampled according to a probability distribution, is grown according to this stochastic rule and also limited in its total number of nodes.

A key statistic we will compute for each use of the Galton-Watson tree null model is the logarithm of the number of equivalent trees for a dendritic arbor,

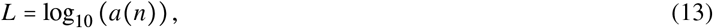

where is the total number of nodes and *a*(*n*) is the number of weakly binary trees that have that have nodes. If we consider a uniform distribution on such trees, then 1 /*a* (*n*) is the probability assigned to each.

Let *p*_branch_ represent the pooled fraction of nodes that have two outgoing edges, and *p*_leaf_ represent the pooled fraction of leaf nodes having 0 outgoing edges. Then, the probability of having a single outgoing edge is 1 − *p*_branch_ −*p*_leaf_. A random Galton-Watson tree, where the number of children at a node is randomly sampled, is grown according to a stochastic algorithm with parameters *p*_branch_ and *p*_leaf_. This simulation is limited in the number of nodes by the number of simulation dendrite nodes. Each node has either two children according to probability *p*_branch_, one child according to probability 1 − *p*_branch_ − *p*_leaf_ or no children according to probability *p*_leaf_.

From a Galton-Watson simulated tree of size *n*, the *n*^th^ Wedderburn-Etherington number counts the number of possible weakly binary trees for *n* nodes. In panel (C) of Fig. VII, the Wedderburn-Etherington number applies to our case because it is a comparison that takes into account runs of no-branch decisions that interleave branch points as in a dendrite arbor tree. Random trees are generated according to the pooled fraction of junction nodes as the branching probability *p*_branch_.

Another key statistic is defined as the histogram of no-branch run lengths. This statistic is shown in Fig. VII Panel D histogram of distances. The histograms are calculated for simulated dendrite trees and Galton-Watson null model trees.

### Statistical analysis

A two one-sided t-tests (TOST) equivalence procedure was used to compare distributions and test for equivalence between simulation and biological dendrite arbor features (20). An initial significance threshold of 0.05 was used, then adjusted using the Holm-Bonferroni correction for multiple testing correction in R. The parameter Cohen’s *d* =1 was chosen for TOST allowing for a large effects margin of one standard deviation. Statistical inference of effect size in biological function research has been studied about Cohen’s *d*. It was estimated for endothelial function that *d* = 1.21 represented a large effect size margin (21). We use a more conservative value of *d* = 1.

### Software and computational details

The version of Mathematica used for simulation is 14.2.1, and Plenum version is 22. The hardware used for runs is AMD EPYC 7H12 64-Core Processor with a CPU in parallel per run. The code written in Mathematica and using Plenum is stored at the GitHub repository: https://github.com/matthewhur836/ClassIDendrite-DGG.

## RESULTS

### Simulated ddaE dendrites recapitulate key morphological features of biological dendrites

We developed and validated a computational model of ddaE neuron dendrite development in the DGG modeling language. This model has a few main components: initial conditions specifying two primary branches directed dorsal and ventral-posterior primary branches; branch growth and shrink velocities; and resource constraints. Two examples of arbors generated by this computational model are shown in Fig. III. Representative images of biological ddaE neurons show striking morphological similarities to our simulated arbors Fig. IV. The model successfully captures key architectural features observed in biological ddaE dendrite arbors.

**Figure 3:**
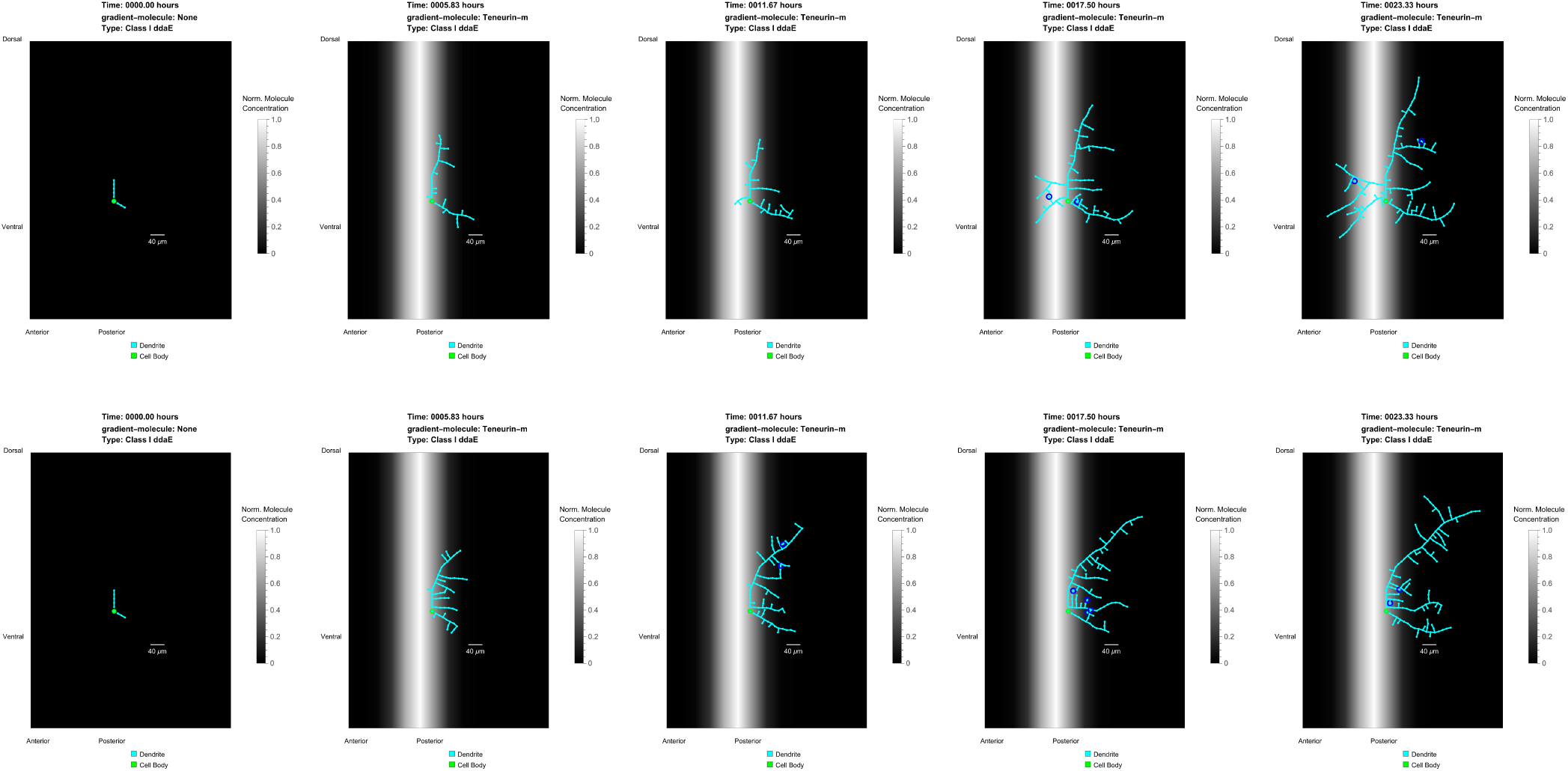
Snapshots of two stochastic simulations of the ddaE class I neuron from the initial condition at t = 0 hours up to t = 23.33 hours. Dark blue circles indicate nodes undergoing self-avoidance retraction. White and grey gradient visualizes the location of the Ten-m gradient during simulation. The cell body is shown as a green circle near the center of each image.

First, simulated arbors begin with a long dorsal primary dendrite growing directly upward from the cell body, and a ventral primary dendrite of variable length, consistent with the primary branch orientations and angles observed in biological ddaE morphology.

Second, secondary and higher-order branches grow asymmetrically from the dorsal and ventral-posterior primary dendrites, with a strong posterior bias. The antero-posterior asymmetry of branches extending from the dorsal primary branch is biased towards the posterior direction, driven by the Ten-m gradient (indicated by the brighter intensity in Fig. III). This computational posterior bias recapitulates the posterior bias characteristic of ddaE neurons.

Third, the resource allocation in both arbors separately constrains the growth to the average total dendritic arbor length. In Fig. III, the difference between snapshots at 23.33 hours and 17.5 hours and between snapshots at 17.5 hours and 11.67 hours in both simulations, upon visual inspection, shows a slowing in the increase of the number of branches. Branches are discouraged to grow later in simulation time because of the implemented resource constraints.

Fourth, dendrite branches do not cross over one another in either Fig. III simulation images or in the set of five snapshots at t = 23.33 hours shown in Fig. IV, matching the self-avoidance behavior of biological dendrites. This maintenance of self-avoidance is consistent with the biological behavior mediated by DSCAM.

**Figure 4:**
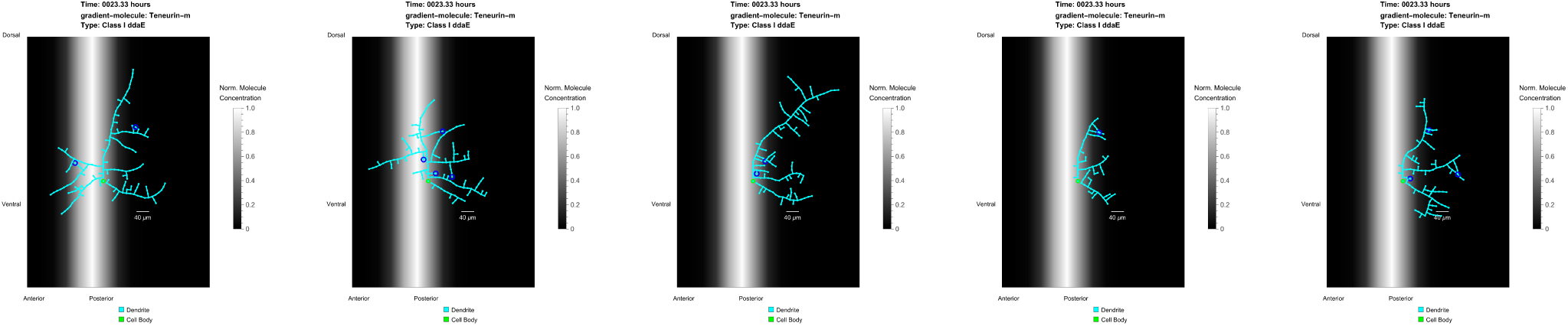
Snapshots of five stochastic simulations of the ddaE class I neuron taken at t = 23.33 hours. Dark blue circle indicates node undergoing self-avoidance retraction. White and grey gradient visualizes the location of the Ten-m gradient during simulation. The cell body is shown as a green circle near the center of each image.

This qualitative similarity between simulated dendrites and biological dendrites is supported with quantitative comparison, discussed below.

### DGG simulations exhibit statistical distributions matching biological neurons

Simulated ddaE arbors closely reproduced the statistical distributions of biological neurons across number of branches, average branch length, and total branch length in both top (dorsal) and bottom (ventral-lateral) arbors (Fig. V; Fig. VI; Table II).

**Table II:**
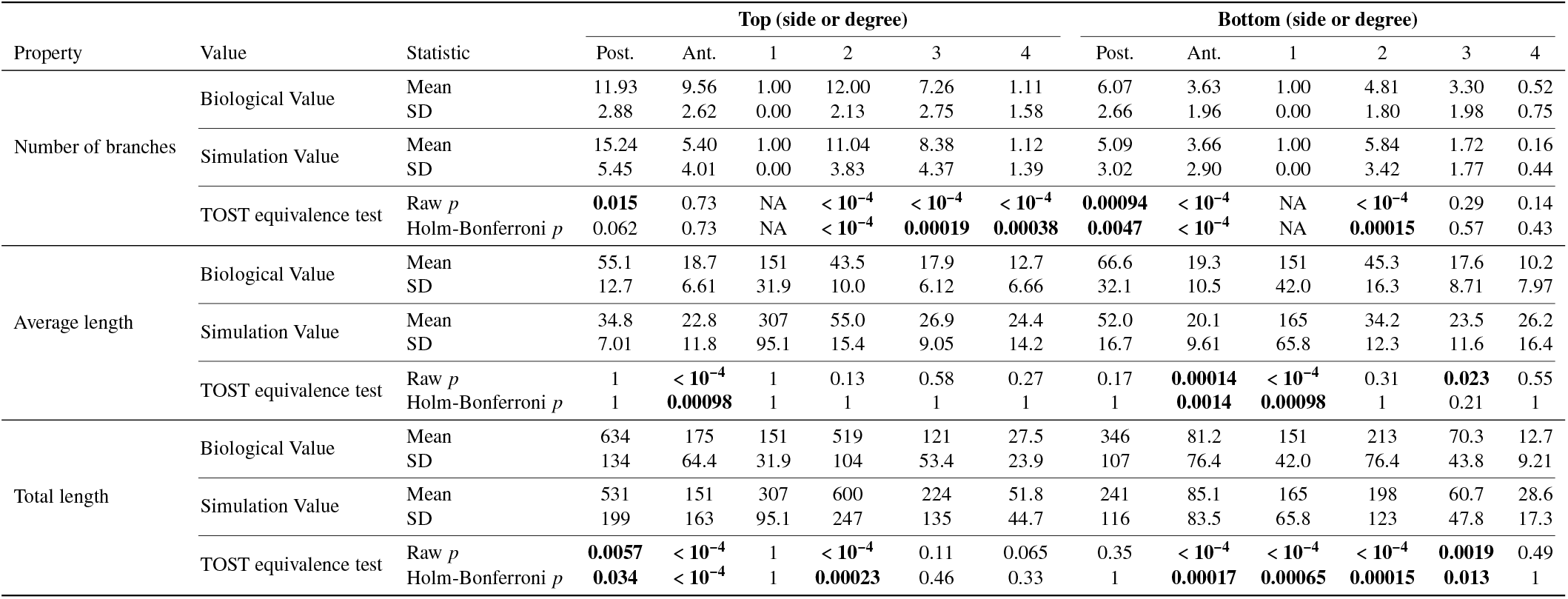
Comparison of biological (n=19 or 8) and simulation (n=100) properties for dendrite branch categories. Mean and standard deviations of samples are provided in the table. Simulation sample size is 100 dendrite trees and biological sample size is 19 other than for degree categories of length measures in which case it is 8. To test equivalence between biologically imaged and computationally simulated dendrites, we use the two one-sided t-tests (TOST) procedure (20) The adjusted p-value threshold after Holm-Bonferroni multiple testing correction for p-value on the TOST equivalence test results is shown below the Raw *p* as “Holm-Bonferroni *p*”. The parameter Cohen’s *d* = 1 allowing for a large effects margin of one standard deviation as explained in the Statistical Analysis section was chosen for TOST.

**Figure 5:**
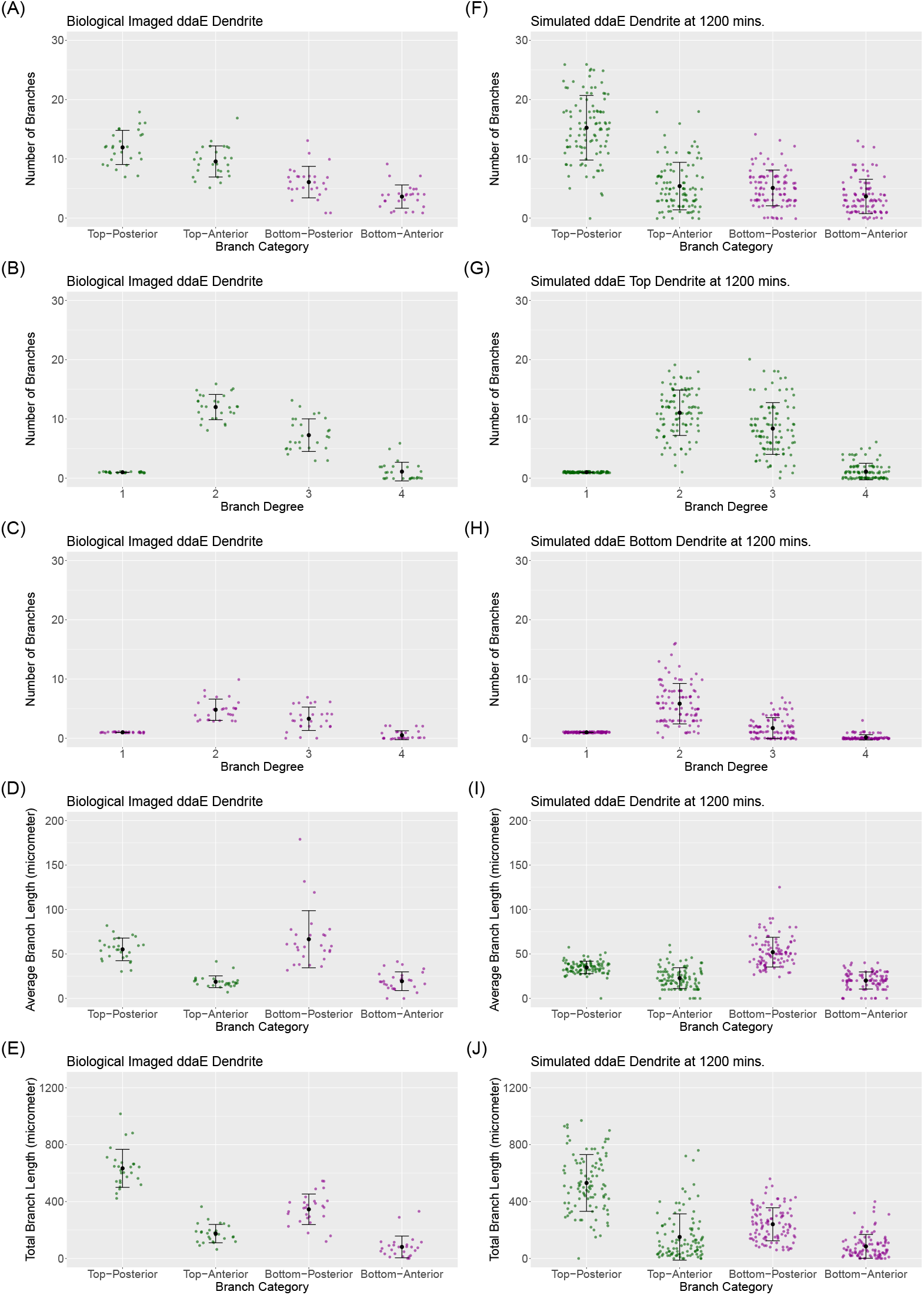
Comparison of characteristic features between biologically imaged dendrites ((A)–(E)) and computationally simulated dendrites ((F)–(J)). (A)–(C) and (F)–(H): distribution of branch counts (from top-to-bottom) (1) based on arbor category, i.e. top (dorsal) vs. bottom (ventral) and anterior vs. posterior, (2) divided by branch degree for the top arbor and (3) divided by branch degree for the bottom arbor. (D)–(E) and (I)–(J): (1) average branch length and (2) total branch length, for branch categories labeled on the horizontal axes (top-posterior, top-anterior, bottom-posterior, bottom-anterior). Black dots represent means and bars represent ± one standard deviations.

**Figure 6:**
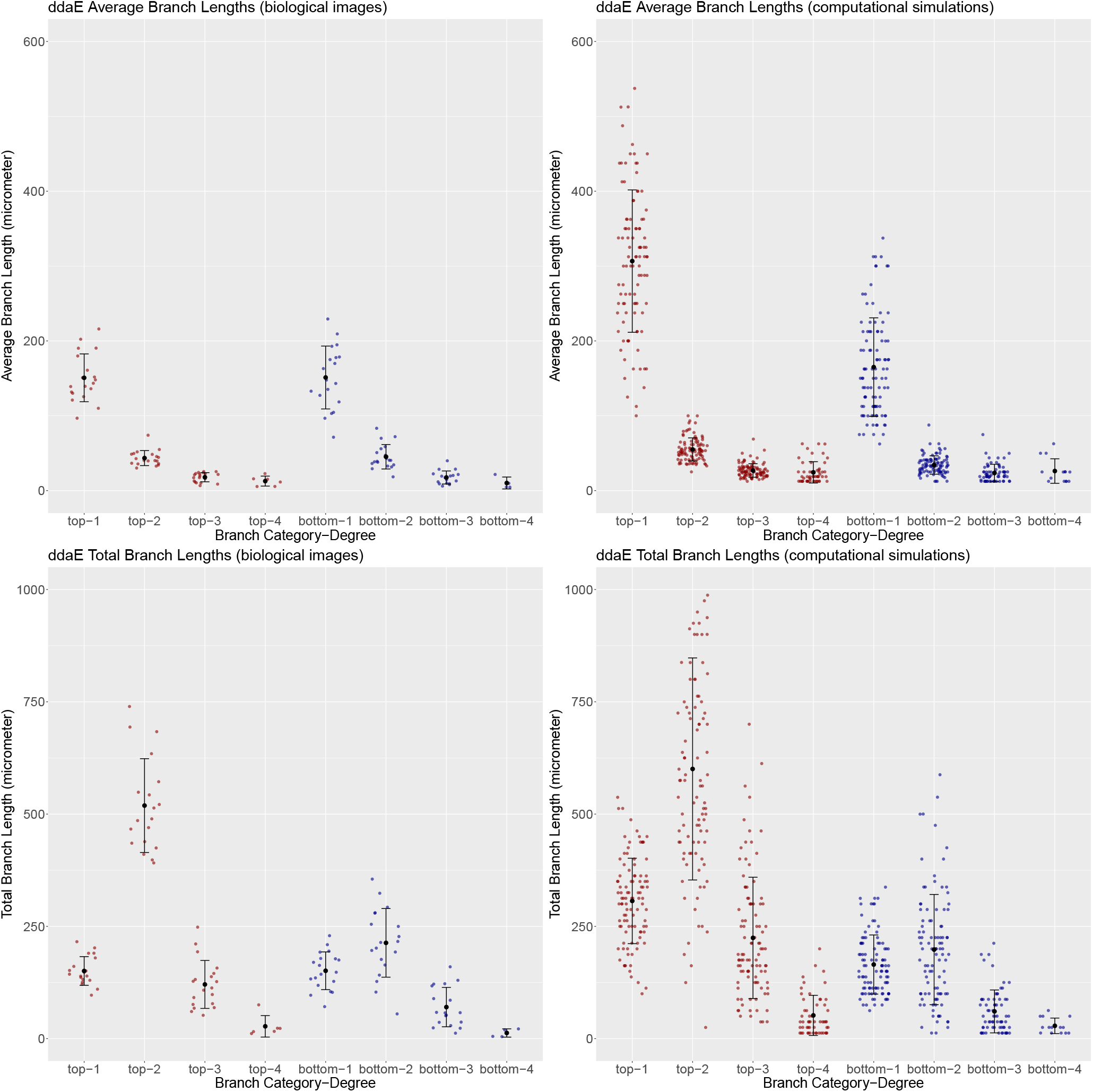
Comparison of characteristic length features, categorized by branch degree, between biologically imaged dendrites and computationally simulated dendrites. Top row: distribution of average branch length (1) based on dendrite arbor type, i.e. top (dark red) or bottom (dark blue) and (2) based on dendrite branch degree (1°, 2 °, 3 °, or 4 °). Bottom row: distribution of total branch length with the same categorization as average branch length. Black dots represent means and bars represent ± one standard deviations.

We quantified the number of branches pointing anteriorly or posteriorly in both the top and bottom arbors in both the biological and the simulated data (Fig. V A, F; Table II). Within both the biological and simulation data, in both the top and bottom arbors, posterior-directed branches outnumbered anterior-directed branches significantly, demonstrating a posterior bias in branch number. In the bottom arbor (purple), the number of posterior-directed branches was equivalent between simulation and biological data (TOST p = 0.0047), as was the number of anterior-directed branches (TOST p < 10^−4^). In the top arbor (green), the number of posterior-directed branches did not achieve equivalence between biology and simulation (TOST p = 0.062); similarly, the number of anterior-directed branches in the top arbor was not significantly equivalent in biological data compared withsimulation (TOST p = 0.73).

We next quantified the number of branches by branch degree in both the top and bottom arbors (Fig. V B, C, G, H; Table II), where degree 1 denotes primary branches attached directly to the soma, and degree 4 denotes quaternary branches. In the top arbor (B, G, green), the number of branches at degrees 2, 3, and 4 were all equivalent between simulation and biological data (TOST p < 10^−4^, 0.00019, and 0.00038, respectively) indicating that the model accurately reproduces branching complexity in the top arbor. In the bottom arbor (C, H, purple), the number of branches at degree 2 were equivalent between simulation and biological data (TOST p < 0.00015). Degrees 3 and 4 were underrepresented in the simulation relative to biology, but absolute branch counts at degrees 3 and 4 are low in both datasets, limiting the concern of this discrepancy.

We next compared average branch length between simulation and biological data (Fig. V D, I; Table II). We observed a bias towards posterior-directed growth in both the biological data and the simulation data, in both the top and bottom arbors, where posterior-directed branches had longer average branch lengths than anterior-directed branches. The posterior bias therefore manifests not only in branch number but also in average branch length, and this directional pattern is reproduced by the model in both top and bottom arbors. Comparing simulation to biological data, average branch length in the top-anterior and bottom-anterior categories was in agreement (TOST p < 0.00098 and 0.0014, respectively). Average branch length in the top-posterior and bottom-posterior was modestly lower in the simulation than in biological data, consistent with the overestimate of posterior branch number seen in the top arbor. When comparing total branch length between simulation and biological data ((Fig. V E, J; Table II), we observed the same patterns: a bias towards posterior growth in both the biological and simulated data, in both the top and bottom arbor; and comparable total branch lengths when comparing biological versus simulated data (Top-posterior TOST p = 0.034, top-anterior TOST p < 10^−4^, and bottom-anterior TOST p = 0.00017). The exception is that bottom-posterior total branch length is not significantly equivalent between simulation and biology (TOST p = 1), with biological values exceeding the simulation.

We compared average branch length by branch degree in both the top and bottom arbors (Fig. V top; Table II). In the top arbor, average branch lengths were longer in the simulation than in biological data for primary, secondary, and tertiary branches, while the biological quaternary branches were not significantly equivalent with the simulation (TOST p = 1), indicating a systematic overestimate of individual branch elongation in the top arbor across most branch degrees. In the bottom arbor, average branch length was equivalent between simulation and biological data for primary branches (TOST p < 0.00098), while secondary, tertiary, and quaternary branches were all longer in the simulation than biological data.

We last compared total branch length by branch degree in both the top and bottom arbors (Fig. V bottom; Table II). In the top arbor, total branch length was in agreement for secondary branches (TOST p = 0.00023), while degrees 1, 3, and 4 were significantly longer in the simulation than in biological data. In the bottom arbor, total branch length was equivalent between simulation and biological data for primary, secondary, and tertiary branches (TOST p = 0.00065, 0.00015, and 0.013, respectively), but not at degree 4, indicating that the overall dendritic investment in the bottom arbor is mostly accurately reproduced by the model.

Together, these analyses demonstrate that the computational model accurately reproduces the key statistical properties of biological ddaE dendrites across both the top and bottom arbors, including number of branches, branch degree distributions, and the directional bias in both average and total branch length. The model reproduces the posterior bias observed in vivo across both branch counts and branch lengths, with the primary systematic discrepancy being an overestimate of posterior growth in the top arbor that is consistent across multiple measures, and a selective underestimate of bottom arbor higher-degree branch counts, which likely reflects a missing anterior-biasing mechanism not captured by the Ten-m gradient alone.

In Table II, the means of posterior and anterior number of branches as well as for primary to quaternary branches are within one simulation standard deviation between simulation and biological values. Under the Holm-Bonferroni multiple testing correction, we find that only the top anterior branches fail to reject the null even by the raw TOST p-value, which is p-value=0.73, with the biological anterior branch count higher than the simulation value. In contrast, the top posterior branches are corrected by Holm-Bonferroni to not reject the null (raw TOST p-value = 0.015, TOST p-value 0.062), which indicates potential for this property to be properly replicated. A number of improvements including increasing sample size to increase statistical power, better inference of the hand-tuned parameters, or parameters substituted by Class I specific biological measurements can all provide avenues for top posterior arbor complexity (number of branches) equivalence test null rejection.

The number of branches subdivide either into posterior and anterior categories or 1° to 4° branch degrees. For our characterization of biologically imaged and computationally simulated dendrites, we notice that in the top row of Fig. V ((A) vs. (F)) that the posterior-anterior bias is somewhat greater in the computationally simulated dendrites than the biologically imaged dendrites. This posterior bias seems however to be in agreement for the ventral-lateral (bottom) arbors between simulation and biological images. Furthermore, characterization of number of branches by branch degree ((B) vs. (G) and (C) vs. (H)) is consistent between simulation and biological images. This comparison suggests that posterior-anterior Ten-m-based bias can more strongly affect the top arbor than the bottom arbor, which follows from the angle of the ventral-lateral away from the Ten-m gradient peak. Furthermore, the simulation branch degree distribution matches the biologically imaged dendrites’ distribution, indicating branch degree can be consistent separate from the strength of posterior bias. Average branch length ((D) vs. (I)) and total branch length ((E) vs. (J)) distributions roughly match between biological images and computational simulations.

### Comparison with null model distributions reveals structural model

To assess whether simulated arbors exhibit non-random structural organization, we use the “dartboard” null model for distance based distributions and the Galton-Watson tree null model for topological properties. We compared the simulated arbors with these null models. For the first null model, the simulation-derived distance between junctions alone, or between all nodes are compared separately using the histogram of Equation 12 in the cases of top and bottom arbors. The distribution of the mean of node-counting distributions is shown in Fig. VII based on the positions of nodes within variable radius *ρ*. There also is such a distance measure between randomly sampled “dartboard points” within each arbor’s bounding box (in red, top panels of Fig. VII).

**Figure 7:**
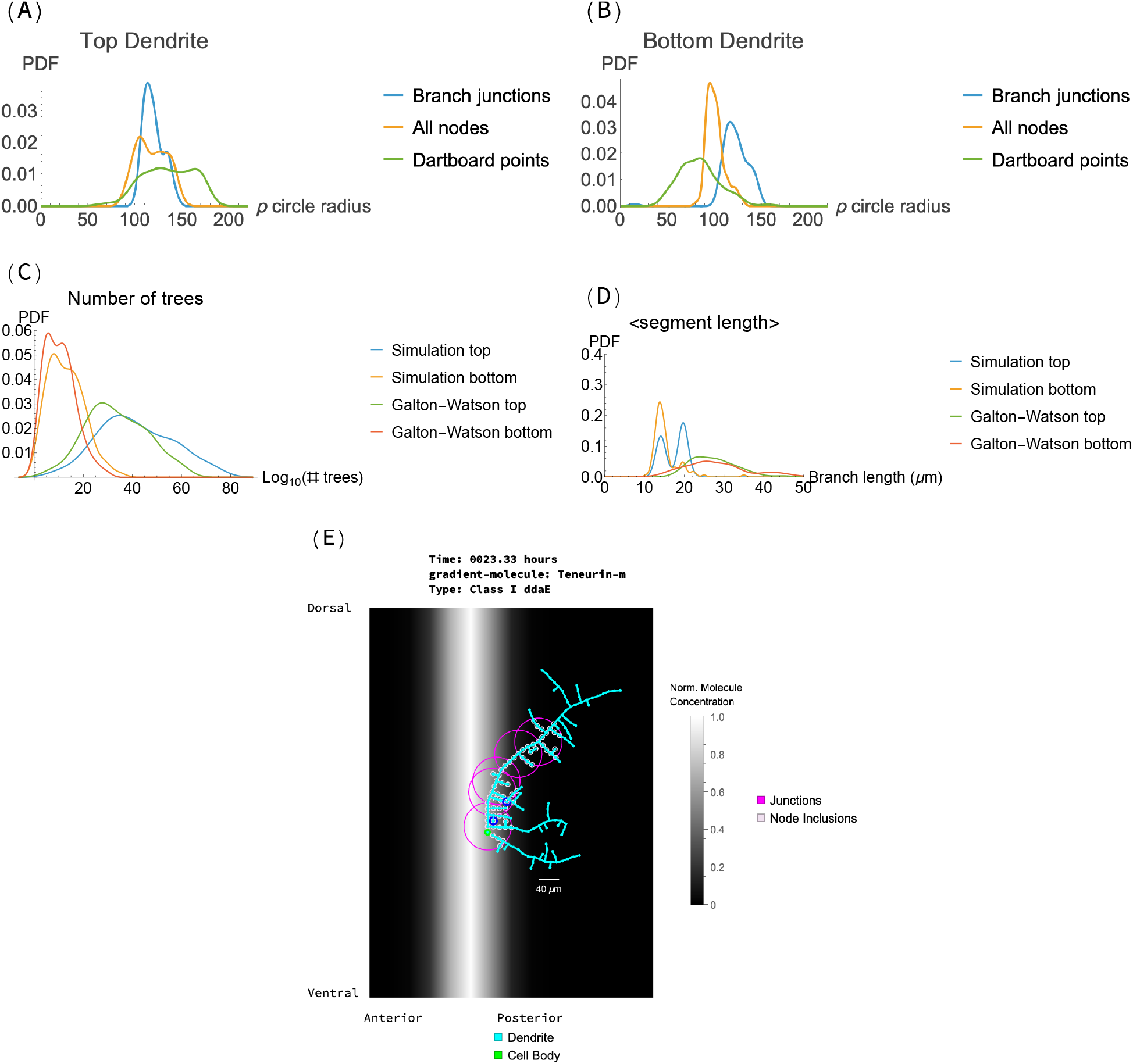
Comparison models for computationally simulated dendrites. (A) node inclusion radius for top dendrites and (B) node inclusion radius for bottom dendrites. (C) number of possible trees with same total node count, (D) average branch segment length (between two junctions and from a junction to a leaf) (E) labeled dendrite arbor for a sample of junction nodes (dark yellow) and node inclusions within radii of *ρ* = 50*μ*m. Panels (A)-(D) show distributions different from the “Dartboard points” null model for panels (A) and (B) and “Galton-Watson tree” null model for panels (C) and (D).

For the Galton-Watson tree null model, the simulation-derived internal node counts are permuted for binary tree structure, by analogy with Equation 13, to yield the total number of possible trees (Fig. VII Panel (C)). Such an enumeration for the null distribution can be efficiently calculated using the count of the number of weakly binary trees (Wedderburn-Etherington) calculatable by their recurrence relation. For the third null model the average segment length, which is a run of no-branch decisions between junction nodes or between a junction and a leaf, is calculated for each simulation and Galton-Watson tree (Fig. VII Panel (D)). A representative example of junction and node classifications is shown in Fig VII Panel (E). Histograms of the distributions are shown in Fig. VII.

The distributions observed in Panels (A) and (B) of Fig. VII demonstrate that simulated arbors exhibit non-random spatial and topological organization, consistent with biological structural constraints. Specifically in Panel (A) no-branch decision runs are most similar to all nodes in their inter-nodal distance, and the branch junctions have a similar mean to all nodes. Compared to an expected greater average distance between a pair of junction points for a dendritic arbor, it is actually similar to between a pair of any nearby simulation nodes. For the top arbor the branch junction set is compactly organized. Top arbors are more compactly organized in all three measure types compared to random dartboard points. This difference supports a principle of efficient spatial organization, where branching leads to compact node positions, while non-branching runs are distributed more randomly. For a bottom arbor in Panel (B) the three measure types are less compact than random dartboard points with distances for inclusion of branch junctions in line with expectation that they are distributed farther apart than for a typical pair of nodes. As the “no-branch decisions” and “all nodes” distributions are similar (blue and green in Panels (A) and (B)) the specific location of a branch does not differentiate from selecting any node for this analysis. Information concomitant with inter-junction location may not reside in the number of nearby nodes, but in each outgrowth’s segment length which is a more specific measure to each segment. Next we discuss comparison models for this hypothesis.

In Panel (C) of Fig. VII, we find that the top arbor’s junction organization (blue) differentiates itself from the Galton-Watson null (green) model in position of peaks more than for the bottom arbor tree counts (yellow) and its null model (red). The organization of internal junction nodes drives top arbor uniqueness more so than for the bottom arbor. Yet, as discussed previously, the average node inclusion radius is similar between junctions and any nearby nodes. Panel (D) ascribes for the top arbor the information in its uniqueness to each no-branch segment’s length, where a bimodality (blue in Panel (D)) is different from the unimodal Galton-Watson null model. The larger segment length peak is similar but the top arbor from simulation has a lower peak indicating there is a second process for more frequent branching. This comparison reveals a consistent explanation between panels (C) and (D) for top arbor branching: there is a process for more frequent branching than average branching fraction of nodes.

The bottom arbor (yellow in Panel (D)) lacks the process for less frequent branching, but also is less compactly organized than random dartboard points (in Panel (B)), indicating that the less frequent branching may form closer compactness as in the inter-nodal vertical distances between runs of posterior secondary branches.

## DISCUSSION

We developed and validated a computational model of ddaE proprioceptive neuron development in Drosophila larvae that accurately recapitulates key morphological features of biological dendrites. Our results demonstrate that a single morphogen gradient (Ten-m) combined with resource allocation constraints and self-avoidance rules, is sufficient to generate the characteristic asymmetric branching growth of the ddaE neurons. Critically, we show that simulated and biological arbors are statistically indistinguishable across the majority of measured morphological properties, with only top anterior branch number differing significantly, establishing this model as a validated platform for investigating genetic and molecular perturbations *in silico*. Our results further indicate that the spatial gradient (finite difference) rather than absolute Ten-m concentration is the critical parameter for directional branch growth, that a single morphogen cue combined with resource constraints and self-avoidance is sufficient to specify the major features of a complete arbor pattern, and that resource allocation constraints and self-avoidance together shape final arbor size and morphology.

Our model demonstrates that a single morphogen gradient (Ten-m) alone can guide the asymmetric elaboration of the ddaE dendrite arbor once the primary dendrites are established. Previous research identified the morphogen Wingless1 (Wg1) of the Wnt signaling pathway as important for establishing primary dendrite morphology (3). Our findings are complementary: we show that the morphological features we examined (branch number, branch length, posterior bias, and self-avoidance) can be recapitulated using Ten-m guided growth from initial conditions that specify two primary dendrites separated by a pre-determined angle of about 120 degrees. This suggests that Wg signaling may establish the primary branch angles and orientations, after which Ten-m guides the elaboration and patterning of higher-order branches. The discrepancy in top anterior branch number between simulation and biological data suggests that at least one additional anterior-biasing mechanism may operate in parallel with Ten-m. The specific roles of Wg1 in setting initial conditions, potential interactions between Wg and Ten-m pathways, and the molecular mechanisms linking receptor activation to cytoskeletal dynamics remain important questions that could be addressed by incorporating Wg1-dependent initial condition variability into future models.

Previous computational models of Drosophila sensory neurons have focused on the highly branched class IV neurons (6, 7), actin branch terminal Class III neurons (22), or symmetric class I neurons (8), but asymmetric class I neurons like ddaE have not been computationally modeled. Class IV neurons have highly elaborate, space-filling arbors that tile the body wall, which is a fundamentally different patterning problem from the asymmetric, non-space-filling ddaE arbor. Our work extends computational dendrite modeling to the asymmetric Class I ddaE neuron, with distinct morphological and functional properties. Our approach uses stochastic simulation within the DGG framework, incorporating biological constraints that are appropriate for ddaE: directional morphogen gradients, asymmetric resource allocations, and self-avoidance rules. This suggests that graphbased dynamical systems provide a natural and flexible framework for simulating dendrite morphogenesis, with class-specific behaviors emerging from different rule modules and parameter values rather than fundamentally different algorithms.

To build our model, we either estimated our parameters or made an assumption that they are similar to that in Class IV dendrites (Table I). Yet, a large number of morphological properties match those in biological, imaged dendrites, which we have verified using 100 unique stochastic simulations. The set of velocities and tip state transition rates result in net growth of our dendritic arbor, with resource constraints counteracting. Our main goal was not accuracy in the amount of time to grow the final dendritic arbor, but to reach a state in which biological properties are recapitulated *in silico*. The recapitulation of biological features indicate our parameter set is robust to variations within net effects and is consequently a valid parameter set for purposes of analyzing phenotype of the Class I dendritic arbor.

Using our null models we have posited that the spatial locations of junctions are important for dendrite growth efficiency and that the specific organized junction sequence in a binary tree is important for distinguishing dorsal (top) and ventral (bottom) dendritic arbors from random binary trees. During studies of class I dendrite regeneration, it will be important to ensure the branching mechanisms active during development are sufficiently activated. If branch junctions are the drivers of morphological characteristics of neuronal dendrites, the measures of dendrite branching locations can be important morphological features relevant to dendritic function, e.g. the inter-junction segment length can be independent to the branching probability.

Class I ddaE neurons function as proprioceptors that detect body curvature during forward locomotion. Our validated computational model provides a tool for exploring how dendritic morphology contributes to proprioceptive function. For example, the posterior bias in branching may optimize coverage of body regions that undergo maximal curvature during forward locomotion. Future work could couple our morphological model with biophysical simulations of mechanotransduction to test how arbor architecture affects sensory encoding.

Beyond developmental questions, our model has potential applications for studying dendrite regeneration. Injury to PNS neurons leads to impaired dendrite regeneration and incomplete functional recovery (23). A calibrated computational model with modifiable parameters enables *in silico* testing of interventions, such as altered morphogen expression, modified adhesion molecule levels, or altered resource allocations, that might improve regeneration outcomes. While our current model simulates normal development, extending it to incorporate injury responses and regeneration dynamics could guide efforts to aid development of therapeutic targets for human interventions.

## AUTHOR CONTRIBUTIONS

Conceptualization, EDM and KTP; methodology, MH, EDM, and KTP; investigation, MH and PTH; writing—original draft, MH and KTP; writing—review & editing, all authors; funding acquisition, EDM and KTP; resources, EDM and KTP; supervision, EDM and KTP.

## DATA AVAILABILITY

Simulation dataset is deposited at the GitHub repository (https://github.com/matthewhur836/ClassIDendrite-DGG).

## DECLARATION OF INTERESTS

The authors declare no competing interests.

## ACKNOWLEDGMENTS

This work was funded in part by U.S. NIH/NIDA Brain Initiative Grant 1RF1DA055668-01 and the Human Frontiers Science Program Grant HFSP—RGP0023/2018 (to EDM), and R00NS097627 from the NIH National Institute of Neurological Disorders and Stroke (to KTP). This work was supported in part by the UC Southern California Hub, with funding from the UC National Laboratories division of the University of California Office of the President (to EDM). KTP is a fellow of the Hellman Family foundation and the Rose Hills foundation. This study was made possible in part through access to the Optical Biology Core Facility of the Developmental Biology Center, a shared resource supported by the Cancer Center Support Grant (CA-62203) and NIH-S10OD032327. The authors thank all members of the lab for their support.

## Notes

https://github.com/matthewhur836/ClassIDendrite-DGG

### Competing Interest Statement

The authors have declared no competing interest.

